# Ixochymostatin, a trypsin inhibitor-like (TIL) protein from *Ixodes scapularis*, inhibits chymase and impairs vascular permeability

**DOI:** 10.1101/2024.10.04.616518

**Authors:** Larissa Almeida Martins, Markus Berger, Jan Kotál, Stephen Lu, Lucas C. Sousa-Paula, Brian J. Smith, Yixiang Zhang, John F. Andersen, Lucas Tirloni

**Affiliations:** Tick-Pathogen Transmission Unit, Laboratory of Bacteriology, Division of Intramural Research, National Institute of Allergy and Infectious Diseases, Hamilton, MT, USA; Department of Biochemistry and Molecular Biology, University of Nevada, Reno, NV, USA; Centro de Pesquisa Experimental, Hospital de Clínicas de Porto Alegre, Porto Alegre, RS, Brazil; Department of Medical Biology, Faculty of Science, University of South Bohemia in České Budějovice, Budweis, Czech Republic; Rocky Mountain Veterinary Branch, Rocky Mountain Laboratories, Division of Intramural Research, National Institute of Allergy and Infectious Diseases, National Institutes of Health, Hamilton, MT, USA; Protein Chemistry Section, Rocky Mountain Laboratories, Division of Intramural Research, National Institute of Allergy and Infectious Diseases, National Institutes of Health, Hamilton, MT, USA; Vector Biology Section, Laboratory of Malaria and Vector Research, Division of Intramural Research, National Institute of Allergy and Infectious Diseases, Rockville, MD, USA

**Keywords:** *Ixodes scapularis*, tick saliva, chymase, inhibitor, trypsin inhibitor-like

## Abstract

Ticks, as pool feeders, obtain a blood meal by lacerating small blood vessels and ingesting the blood that flows to the feeding site, which triggers various host-derived responses. However, ticks face the challenge of wound healing, a process involving hemostasis, inflammation, cell proliferation and migration, and remodeling, hindering blood acquisition. To overcome these obstacles, tick salivary glands produce a diverse array of bioactive molecules. Here, we characterize ixochymostatin, an *Ixodes scapularis* protein belonging to the trypsin inhibitor-like (TIL) family. It is expressed in multiple developmental stages and in tick salivary glands and acts as a slow and tight-binding inhibitor of chymase, cathepsin G, and chymotrypsin. Predictions for the tertiary structure complex between ixochymostatin and chymase suggest a direct interaction between the inhibitor’s reactive site loop and protease active sites. *In vitro*, ixochymostatin protects the endothelial cell barrier against chymase degrading action, decreasing cell permeability. *In vivo*, it reduces vascular permeability induced by chymase and compound 48/80, a mast cell degranulator agonist, in a mouse model. Additionally, ixochymostatin inhibits the chymase-dependent generation of vasoconstrictor peptides. Antibodies against ixochymostatin neutralize its inhibitory properties, with epitope mapping identifying potential neutralization regions. Ixochymostatin emerges as a novel tick protein modulating host responses against tick feeding, facilitating blood acquisition.

**Highlights:**

- Ixochymostatin is a newly described trypsin inhibitor-like (TIL) protein from *Ixodes scapularis*.
- Ixochymostatin inhibits chymase and reduces vascular permeability.
- Ixochymostatin inhibits chymase-dependent generation of vasoconstrictive peptides.
- Antibodies generated against ixochymostatin neutralize its inhibitory functions.
- Potential epitopes responsible for anti-ixochymostatin neutralization were mapped.

## 1. Introduction

The tick *Ixodes scapularis*, also known as the black-legged tick or the deer tick, is widely distributed in the east part of the United States, where is the most important vector of *Borrelia burgdorferi*, the causative agent of Lyme disease [1]. Ticks are pool feeders that obtain a blood meal by lacerating small blood vessels and ingesting the blood that flows to the feeding site [2]. This feeding mechanism elicits various responses from the host, prompting ticks to confront the challenge of wound healing. Wound healing is accomplished through four precisely orchestrated phases: hemostasis, inflammation, cell proliferation and migration, and remodeling [3]. Collectively, these phases serve as obstacles to the acquisition of a blood meal [4]. In response to these barriers, tick salivary glands have evolved a sophisticated and intricate pharmacological arsenal comprising bioactive molecules aimed at facilitating blood feeding [5]. Tick saliva contains hundreds of compounds that hinder wound healing, possessing anti-coagulant, vasodilatory, anti-inflammatory, and immunomodulatory properties [2]. While aiding the vector in obtaining a blood meal, saliva also alters the site where pathogens are injected and often facilitates the infection process [6].

The pplication of transcriptomics and proteomics has facilitated the identification of a large set of tick salivary molecules that impact tick feeding and play a role in the transmission of tick-borne pathogens. Several tick sialomes studies have been performed and reported that ticks express different classes of proteins [5,7]. Among these classes of salivary proteins, protease inhibitors are highly abundant. Most of the salivary protease inhibitors described belongs to the Kunitz-type, serpin, cystatin, and trypsin inhibitor-like (TIL) family of protease inhibitors [5,7]. Several protease inhibitors have been functionally characterized and shown to impair wound healing, and most of these inhibitors display unique specificity and high-affinity binding for proteases with a role in this process [8].

Although highly abundant in the saliva and salivary glands of ticks, data on the functional characterization of proteins belonging to the trypsin inhibitor-like (TIL) family are still scarce. A protein containing a TIL domain was first described in nematodes of the genus *Ascaris* as an inhibitor of pancreatic trypsin. These inhibitors were initially classified in the *Ascaris* family of serine protease inhibitors [9,10]. Structure-level characterization was first described for the chymotrypsin and porcine pancreatic elastase inhibitor C/E 1 (chymotrypsin/elastase inhibitor-1) isolated from *Ascaris suum* [11]. The crystal structure of the complex between C/E 1 inhibitor and porcine pancreatic elastase was resolved and it was confirmed that these inhibitors belong to a new family of protease inhibitors [12], nowadays being classified in the inhibitor family I8 of MEROPS [13]. Since then, proteins containing the TIL domain have been described and characterized in invertebrates as nematodes of the genus *Ascaris* and *Ancylostoma* [11,14], in the hemolymph of the *Apis mellifera* bee [15], and in vertebrate animals, as the trypsin and thrombin inhibitor present in the skin secretion of the frog *Bombina bombina* [16]. In hematophagous arthropods, only a small number of proteins containing the TIL domain have been identified and characterized. The first inhibitor described was ixodidin, which was initially described as an antimicrobial peptide isolated from the hemolymph of the cattle tick *Rhipicephalus microplus*. Ixodidin inhibits elastase and chymotrypsin with inhibition constants in the nanomolar range [17]. The second inhibitor containing the TIL domain to be described in hematophagous arthropods was BmSI-7, an inhibitor purified from eggs of the tick *R. microplus*. BmSI-7 inhibits neutrophil elastase, subtilisin A, and Pr1 proteases from the entomopathogenic fungus *Metarhizium anisopliae* with inhibition constants in the nanomolar range [18].

Here, we present the first functionally characterized TIL from *I. scapularis*. Ixochymostatin inhibits chymase, cathepsin G, and chymotrypsin. It was identified as a slow and tight-binding inhibitor of chymotrypsin. Inhibition of chymase effectively impairs endothelial cell and vascular permeability and the generation of vasoconstrictor peptides promoted by this protease. Consequently, ixochymostatin emerges as a novel tick protein modulating vascular functions.

## 2. Materials and Methods

### 2.1. Ethics statement

Animal experiments were conducted in accordance with the guidelines of the National Institutes of Health on protocols approved by the Rocky Mountain Laboratories Animal Care and Use Committee. The Rocky Mountain Veterinary Branch is accredited by the International Association for Assessment and Accreditation of Laboratory Animal Care (AAALAC).

### 2.2. Bioinformatic analysis

The sequence of the ixochymostatin (ISCW002768-RA) was downloaded from NCBI. The presence and location of a signal peptide was predicted using SignalP v. 6.0 [19]. The signal peptide was removed from the protein sequence, and the prediction of tridimensional structures was performed using AlphaFold2 [20], which is available on the NIH HPC Biowulf cluster (http://hpc.nih.gov). The full-length mature monomeric structure of ixochymostatin was generated using the standard AlphaFold-2 build, while the ixochymostatin-chymase complexes were constructed using the multimer extension. Five models were generated for each prediction, and the model with the highest pLDDT (predicted local distance difference test) score was selected. The structures were visualized, and the final images were created using the PyMOL Molecular Graphics System (The PyMOL Molecular Graphics System, Version 2.6.0a0, Schrödinger, LLC.).

The deduced mature amino acid sequence of ixochymostatin was used to identify the best match hits in other organisms using BLASTp against the NR and TSA-NR databases from the NCBI. The best match hit for each species was selected, and all sequences were scanned and trimmed for the signal peptide sequence using SignalP v. 6.0 [19]. Amino acid sequences were aligned using the MAFFT algorithm [21]. For phylogenetic analysis, the best-fit evolution model was selected using ModelFinder [22] implemented by IQ-TREE 2 [23] and chosen according to Bayesian information criterion (BIC). The maximum-likelihood phylogenetic reconstruction was inferred by using the IQ-TREE 2 with node support values calculated using 1,000 ultra-fast bootstraps. The phylogenetic tree was plotted and edited using iTOL v5 [24].

### 2.3. Tick feeding, sample collection, and transcription analyses

The *Ixodes scapularis* utilized in this study were purchased from the tick laboratory at Oklahoma State University (Stillwater, OK, USA). Up to 100 larvae, 50 nymphs, and a maximum of 20 adult female ticks were placed on the outer portion of the New Zealand rabbit ear for feeding. All groups were collected in triplicate. Two biological sample groups were generated for larvae: unfed and fully fed, with pools composed of 50 larvae each. The nymph groups consisted of ten individuals each: unfed, as well as at 12, 24, 36, 48, 72, and 96 hours of feeding. For adults, six groups were collected during the feeding stage, along with one unfed group. A pool of five females was manually detached based on their approximate weight ranges (G1: 2.5-3 mg, G2: 6-7 mg, G3: 15-20 mg, G4: 25-30 mg, G5: 45-50 mg, and G6: 105-120 mg) [25]. Within the first hour of detachment, all tick groups were washed, and larvae, nymphs, and adult female salivary glands (SG) and midguts (MG) were homogenized in RNAlater® solution (Thermo Fisher Scientific, Waltham, MA, USA) and stored for later use.

All the samples underwent isolation of total RNA using the AllPrep DNA/RNA/Protein Mini Kit (QIAGEN, Germantown, MD, USA) following the manufacturer’s specifications. Five hundred nanograms of total RNA were treated with DNase I (Thermo Fisher Scientific, Waltham, MA, USA) and reverse transcribed (RT) into cDNA using M-MLV Reverse Transcriptase (Thermo Fisher Scientific, Waltham, MA, USA) according to the manufacturer’s instructions. The cDNA was quantified, and standard curves were generated using different concentrations of cDNA (ranging from 6.25 to 400 ng) to determine the efficiency of each pair of primers. Ixochymostatin specific primers (Forward: 5′-CGATTGGGACTGCGTATCAC −3′ and Reverse: 5′-CTCGATTCCTGTGTTGGCTG-3′) and the ribosomal protein IsRpL13a was used as reference [26] (Forward: 5′-CCTGGACAGGCTCAAGATGT −3′ and Reverse: 5′-TTCTGGTACTTCCAGCCCAC-3′). These primers were synthesized by Eurofins Genomics (Louisville, KY, USA). The qPCR was performed using 100 ng of cDNA as a template with SYBR® Green Master Mix on a CFX96 Real-Time PCR System (Bio-Rad Laboratories, Hercules, CA, USA), following the program: 95°C for 10 min followed by 40 cycles at 95°C for 15 s, 60°C for 60 s, and 72°C for 20 s. Relative quantification of the ixochymostatin transcript was determined using the deltaCt method [27]. Each biological replicate was analyzed in a technical triplicate.

### 2.4. Ixochymostatin recombinant expression and purification

The protein-coding sequence of ixochymostatin was codon-optimized for mammalian expression and synthesized by BioBasic Inc. (Markham, ON, Canada). The mature protein-coding sequence was then cloned into the VR2001 vector [28] in frame with the tPA signal peptide sequence, containing a 6x-histidine tag followed by an enterokinase cleavage site. Plasmid DNA was purified using NucleoBond PC 2000 plasmid megaprep kits (Takara, San Jose, CA, USA). FreeStyle 293-F and Expi293 human embryonic kidney cells (Thermo Fisher Scientific, Waltham, MA, USA) were transfected with endotoxin-free plasmid DNA at the Protein Expression Laboratory of Leidos Biomedical Research, Inc. (Frederick, MD, USA). Supernatants were collected 72 hours post-transfection, centrifuged at 1,000 g for 15 minutes, frozen, and shipped to our laboratory. The recombinant protein was purified through a series of chromatographic steps including affinity, ion exchange, and size-exclusion chromatography. Initially, the supernatant containing the secreted recombinant protein expressed using FreeStyle 293-F cells was applied to a HisTrap excel column (Cytiva, Marlborough, MA, USA) using a peristaltic pump. The column was then washed with 2 column volumes of Tris-HCl 20 mM, NaCl 150 mM, pH 7.4 buffer, and the recombinant protein was eluted using a linear gradient of Tris-HCl 20 mM, NaCl 150 mM, imidazole 500 mM, pH 7.4 buffer on an ÄKTA start system (Cytiva, Marlborough, MA, USA). Subsequently, the eluted fractions were pooled, concentrated, and dialyzed against Tris-HCl 20 mM, pH 8.0 using an Amicon® Ultra-15 centrifugal filter unit (Millipore Sigma, Burlington, MA, USA) for ion exchange chromatography. The sample was then applied to a pre-equilibrated HiTrap Q column, washed with 5 column volumes of Tris-HCl 20 mM, pH 8.0 buffer, and eluted using a linear gradient of Tris-HCl 20 mM, NaCl 1 M, pH 8.0 buffer on an AKTA high-performance liquid chromatography system (Cytiva, Marlborough, MA, USA). Following ion exchange chromatography, the eluted fractions were pooled, concentrated, and dialyzed against Tris-HCl 20 mM, NaCl 150 mM, pH 7.4 buffer using Amicon® Ultra-15 centrifugal filter units (Millipore Sigma, Burlington, MA, USA) before undergoing gel filtration chromatography. The sample was loaded onto a Superdex 75 10/300 GL column, and proteins were eluted isocratically at a flow rate of 0.5 mL/minute in Tris-HCl 20 mM, NaCl 150 mM, pH 7.4 buffer. Eluted protein fractions were pooled, and the protein concentration was determined using the BCA assay (Thermo Fisher Scientific, Waltham, MA, USA). The purified protein was then stored at −80°C until use. Throughout the purification process, all protein fractions were visualized via 4-20% SDS-PAGE gel electrophoresis, followed by Coomassie blue staining.

Intact mass analysis of purified recombinant ixochymostatin was analyzed by mass spectrometry (MS). Recombinant protein was desalted using SP3 protocol [29] and concentrated to 10 pmol/µL solution in 50% acetonitrile (ACN) and 0.1% trifluoroacetic acid (TFA). Intact protein analysis was carried out using Q Exactive plus (Thermo Fisher Scientific, Waltham, MA, USA) mass spectrometer. Samples were directly infused using a syringe pump (Thermo Fisher Scientific, Waltham, MA, USA) with a flow rate of 5 μL/min. The instrument was operated at a resolution of 280k, spray voltage of 3.5 kV, capillary temperature of 275 °C, 4 microscans, and a mass range 1000-3000 m/z. The Xtract algorithm of the Freestyle (version 1.5) was used to deconvolute the raw data. Deconvolution parameters were set as Output Mass, MH+; Charge Range, 5 to 20; Min Num Detected Charge, 3.

### 2.5. Serine protease inhibition screening

Human thrombin (2 nM), human factor Xa (2 nM), human plasmin (20 nM), human kallikrein (2 nM), human factor XIa (5 nM), human factor XIIa alpha (20 nM), and human factor XIIa beta (20 nM) were obtained from Enzyme Research Laboratories (South Bend, IN, USA). Human chymase (20 nM), bovine pancreatic chymotrypsin (5 nM), and porcine pancreatic trypsin (1 nM) were obtained from Sigma-Aldrich (St. Louis, MO, USA). Human cathepsin G (380 nM), porcine pancreatic elastase (50 nM), human neutrophil elastase (25 nM), human uPA (20 nM), and human tPA (20 nM) were obtained from Innovative Research, Inc. (Novi, MI, USA). Substrates were used at 200 μM final concentration: H-D-Phe-L-Pip-Arg-pNA (S-2238) for thrombin; Bz-Ile-Glu(γ-OR)-Gly-Arg-pNA (S-2222) for factor Xa; H-D-Pro-Phe-Arg-pNA (S-2302) for factor XIa, factor XIIa alpha, factor XIIa beta, and human kallikrein; H-D-Val-Leu-Lys-pNA (S-2251) for plasmin; H-D-lle-Pro-Arg-pNA (S-2288) for trypsin, tPA, and uPA (Diapharma Inc., West Chester, OH, USA). Substrates N-Succinyl-Ala-Ala-Ala-pNA (S4760) for pancreatic elastase; N-Succinyl-Ala-Ala-Pro-Phe-pNA (S7388) for cathepsin G, chymotrypsin and chymase (Sigma Aldrich, St. Louis, MO, USA); and MeOSuc-Ala-Ala-Pro-Val-pNA (324696) for neutrophil elastase (Millipore Sigma, Burlington, MA, USA).

All assays were performed at 30 °C in triplicate using 96-well plates. Ixochymostatin (1 μM) was pre-incubated with each protease for 10 minutes in 20 mM Tris-HCl, 150 mM NaCl, 0.01% Tween-20, pH 8. After incubation, the corresponding substrate for each protease was added to a 100 μL final reaction volume. The substrate hydrolysis rate was followed at OD_405nm_ in kinetic mode using a Synergy H1 plate reader (Agilent, Santa Clara, CA, USA) for 15 minutes with reading intervals of 15 seconds. The observed substrate hydrolysis rate in the absence of ixochymostatin was considered 100% and compared with the residual enzymatic activity in the presence of the inhibitor. Data are presented as mean ± standard error of triplicate readings.

### 2.6. Kinetics studies

To determine whether ixochymostatin exhibits a slow- or fast-binding mode, progress curves were generated with and without a 10-minute preincubation of chymotrypsin (5 nM) and ixochymostatin (50 nM). Kinetics assays were conducted using the chromogenic substrate N-Succinyl-Ala-Ala-Pro-Phe-pNA on a Synergy H1 plate reader to assess the type of inhibition, binding mode, and kinetic constants, including half-maximal inhibitory concentration (IC_50_) and apparent Ki (*K_i_^app^*). All assays were performed at 30°C in 100 µl of 20 mM Tris-HCl, 150 mM NaCl, and 0.01% Tween-20 at pH 8. Ixochymostatin (0 – 50 nM) was incubated with chymotrypsin (5, 15, and 25 nM) at 30°C for 10 minutes. Following this incubation substrate was added, and the reactions monitored for 15 minutes. The rate of substrate hydrolysis observed in the absence of ixochymostatin was considered 100% serving as a baseline for comparison against the residual enzymatic activity in the presente of the inhibitor. A positive linear regression was used to plot the IC_50_ values against the protease concentrations to identify the tight-binding mode of inhibition. Additionally, the *K_i_ ^app^* was determined by incubating chymotrypsin (25 nM) with increasing concentrations of ixochymostatin (0 – 96 nM) in 100 µl of 50 mM Tris-HCl, 150 mM NaCl, and 0.01% Tween-20 at pH 8.0 for 10 min at 30 °C. After this incubation, N-Succinyl-Ala-Ala-Pro-Phe-pNA (200 µM) was added, and the reaction was followed by 15 minutes. The inhibition curve was performed in three technical replicates, and the average residual activity was plotted as a function of ixochymostatin concentration. The Morrison’s equation [30] was fitted to the data using the GraphPad Prism software (GraphPad Software, Boston, MA, USA).

### 2.7. Surface plasmon resonance (SPR) assays

Evaluation of binding kinetics by SPR was performed using a Biacore T200 instrument (Cytiva, Marlborough, MA, USA). Recombinant ixochymostatin was immobilized on a CM5 chip to a level of 975 resonance units (RU) at in 10 mM sodium acetate buffer (pH 5.0) using the amine coupling method. After conditioning, chymotrypsin (0 – 31.2 nM) was passed over the surface and kinetic data were collected in single cycle mode using a running buffer of 10 mM HEPES pH 7.4, 150 mM NaCl (HBS-) at 25°C. The data were fit to a 1:1 Langmuir binding model using the Biacore evaluation software.

### 2.8. Angiotensin II and endothelin I generation assays

Ten micrograms of human angiotensin I (Anaspec, Fremont, CA, USA) were incubated at 37°C for 30 minutes in PBS, pH 7.4, with human chymase (50 nM) and various concentrations of ixochymostatin (12.5 - 300 nM). Negative controls included only angiotensin I (10 µg), and angiotensin I (10 µg) plus ixochymostatin (300 nM), while the positive control comprised angiotensin I (10 µg) plus chymase (50 nM). The same procedure was applied to assess big endothelin (Anaspec, Fremont, CA, USA) cleavage.

After incubation, all samples were processed for mass spectrometry analysis. Samples were desalted using a C18 Zip-tip (Millipore Sigma, Burlington, MA, USA) and dissolved in 200 μL of reconstitution buffer (composed of 50% CAN and 1% FA). A Q Exactive Plus mass spectrometer (Thermo Fisher Scientific, Waltham, MA, USA) was utilized for ESI-MS experiments, operating at a resolution of 280k, spray voltage of 3.5 kV, and capillary temperature of 275°C. Samples were infused directly using a syringe pump (Thermo Fisher Scientific, Waltham, MA, USA) at a flow rate of 5 μL/min. Xtract software was employed for deconvoluting the raw data.

### 2.9. Endothelial cell permeability

Endothelial cell permeability was assessed using the FITC-dextran method with transwell inserts (12 mm × 0.4 μm) (Millicell Cell Culture Insert, Millipore Sigma, Burlington, MA, USA). Human microvascular endothelial cell 1 (HMEC-1) at a density of 5 × 10^4^ cells per transwell insert were seeded onto 1% gelatin-coated inserts (24 wells) and cultured for 7 days until a 100% confluent monolayer was formed within each insert. The inserts were then divided and subjected to the following treatment groups (n = 4 per group per plate): (*i*) PBS, pH 7.4, (*ii*) chymase (50 nM), (*iii*) chymase (50 nM) plus ixochymostatin (250 nM), and (*iv*) ixochymostatin (250 nM). Treatments were applied to the upper chamber in Krebs-Ringer bicarbonate buffer containing 2 g/L BSA and FITC-dextran (70 kDa, 1 mg/mL) in a total volume of 200 μL, while the lower chamber received only Krebs-Ringer buffer in a total volume of 500 μL. FITC-dextran leakage into the lower chamber was measured in duplicate for each insert at 15, 30, 60, 120, and 240 minutes using the Cytation 5 multimode reader (Agilent, Santa Clara, CA, USA) at an excitation wavelength of 485 nm and an emission wavelength of 528 nm. Throughout the experiment, cells were maintained in a 37 °C/5% carbon dioxide environment. Additionally, endothelial cell permeability was evaluated through vascular endothelial-cadherin (VE-cadherin) immunocytochemistry. HMEC-1 cells at 100% confluence were treated with PBS pH 7.4, chymase (50 nM), chymase (50 nM) plus ixochymostatin (250 nM), and ixochymostatin (250 nM). Following a 4-hour incubation at 37°C/5% carbon dioxide, cells were fixed in methanol, permeabilized in Tris-buffered saline/0.1% Tween, blocked in 10% normal goat serum, and subjected to overnight immunolabeling with VE-cadherin antibody (working dilution 5 μg/mL) (Life Technologies, Carlsbad, CA). The secondary antibody used was a 448-fluorescence conjugated goat anti-rabbit IgG at a dilution of 1:1,000 (Life Technologies), and nuclei were counterstained with Hoechst. Fluorescently labeled images were captured using the Cytation 10 confocal imaging reader (Agilent, Santa Clara, CA, USA).

### 2.10. *In vivo* vascular permeability

Vascular permeability was evaluated *in vivo* using BALB/c mice (both male and female) maintained under general inhalation anesthesia with isoflurane vaporized in 100% oxygen. Anesthesia was induced at a dose of 3-5% isoflurane and maintained at 0.5–3% with a flow rate of 0.5–2 L/min throughout the experiment. The dorsal region of the mice was shaved, and then each mouse received a retro-orbital injection in one eye of Evans blue dye (50 mg/kg in PBS pH 7.4, 50 μL per injection). After 5 minutes, intradermal injections were administered on the dorsal region (50 μL per site) with the following treatments: (*i*) PBS pH 7.4, (*ii*) compound 48/80 (1 µg), (*iii*) ixochymostatin (5 µg), (*iv*) chymase (3 µg), (*v*) compound 48/80 (1 µg) plus ixochymostatin (5 µg), and (*vi*) chymase (3 µg) plus ixochymostatin (5 µg). Each treatment was applied twice per animal, resulting in a total of 20 spots per treatment (n = 10 mice). After 1 hour, the mice were euthanized, and the skin encompassing both injection sites was carefully removed and photographed. Evans blue dye spots on the skin were excised, and the dye from each spot was extracted by incubating the skin with 1.5 mL of 50% formamide for 24 hours at 55°C. Following centrifugation at 10,000 rpm for 10 minutes, the absorbance of the supernatant was measured at OD_620nm_.

### 2.11. Monospecific antibody production and neutralization assays

Anti-sera against recombinant ixochymostatin were raised in two rabbits (GenScript, Piscataway, NJ, USA). Total IgG was purified from sera by affinity chromatography (HiTrap™ Protein G HP - Cytiva, Marlborough, MA, USA). Monospecific antibodies were purified as follows. First, 0.35 mg of recombinant ixochymostatin was coupled with 0.1 mg of Sepharose 4B CNBr-activated resin following the manufacturer’s instructions (Sigma-Aldrich, St. Louis, MO, USA). Subsequently, total IgG purified from anti-sera was added to the resin with end-over-end rocking at room temperature for 2 hours. The resin was rinsed with at least 30 mL of PBS, pH 7.4 and the bound monospecific anti-ixochymostatin antibodies were eluted with 100 mM glycine-HCl, pH 2.4 (neutralized with 2 M Tris base, pH 8). Eluted antibody fractions were dialyzed against PBS, pH 7.4 and were analyzed by SDS-PAGE followed by Coomassie blue staining and by immunoblot analysis. Total IgG from pre-immune sera was purified by affinity chromatography (HiTrap™ Protein G HP, Cytiva, Marlborough, MA, USA).

The neutralization of ixochymostatin by monospecific antibodies was determined using a colorimetric enzymatic assay. Ixochymostatin (50 nM) was pre-incubated with monospecific antibodies (varying from 31.25 to 500 nM) for 30 min at 37 °C in a buffer containing 20 mM Tris-HCl, 150 mM NaCl, 0.02% Tween, pH 7.4. This was followed by a second incubation with 10 nM of bovine pancreatic chymotrypsin for 15 min at 37 °C. The chromogenic substrate N-Succinyl-Ala-Ala-Pro-Phe-pNA (final concentration of 0.20 mM) was added to reach a final reaction volume of 100 µL and substrate hydrolysis was measured at OD_405nm_ every 15 s for 15 min at 30°C using the Synergy H1 microplate reader (Agilent, Santa Clara, CA, USA). The observed substrate hydrolysis rate in the absence of ixochymostatin was considered 100% and compared with the remaining enzymatic activity in the presence of the inhibitor. Data are presented as mean ± standard error of triplicate readings.

### 2.12. Epitope mapping of ixochymostatin

The epitope mapping was performed by Pepscan (Pepscan Presto BV, The Netherlands). To reconstruct the epitopes of ixochymostatin, a library of peptide-based mimics was synthesized using Fmoc-based solid-phase peptide synthesis. Synthesis of structural mimics peptides was done using Pepscan’s proprietary chemically linked peptides on scaffolds (CLIPS) technology. The CLIPS technology allows to structure of peptides into single loops, double-loops, triple loops, sheet-like folds, helix-like folds, and combinations thereof [31,32]. In brief, a library of overlapping peptides was synthetized in different sets based on the mature sequence of ixochymostatin. These sets were categorized as follows: (i) LIN9, linear peptides of length 9 residues with an offset of one residue; (ii) LIN12, linear peptides of length 12 residues with an offset of one residue; (iii) LIN15, linear peptides of length 15 residues with an offset of one residue; (iv) LOOP9, constrained peptides of length 9. On positions 2 – 8 are incorporated 7-mer peptides derived from the target sequence of ixochymostatin with an offset of one residue. Cysteine residues were inserted on positions 1 and 9 and joined by mP2 CLIPS in order to create a loop mimic. Native cysteines are replaced by cysteine-acm; (v) LOOP12, constrained peptides of length 12. On positions 2 – 11 are incorporated 10-mer peptides derived from the target sequence of ixochymostatin with an offset of one residue. Cysteine residues were inserted on positions 1 and 12 and joined by mP2 CLIPS in order to create a loop mimic. Native cysteines are replaced by cysteine-acm; (vi) LOOP15, constrained peptides of length 15. On positions 2 – 14 are incorporated 13-mer peptides derived from the target sequence of ixochymostatin with an offset of one residue. Cysteine residues were inserted on positions 1 and 12 and joined by mP2 CLIPS in order to create a loop mimic. Native cysteines are replaced by cysteine-acm; (vii) BET22, β-turn peptide mimics of length 22. On positions 2 – 21 are 20-mer peptides derived from the target sequence of ixochymostation with an offset of one residue. Residues on positions 11 and 12 are replaced by “PG” motif in order to induce the β-turn formation. Cysteine residues were inserted on positions 1 and 22 and joined by mP2 CLIPS in order to stabilize the mimic. Native cysteines are replaced by cysteine-acm; and (viii) HEL19, α-helical peptide mimics of length 19 derived from residues of the target sequence with an offset of one residue. Cysteines are inserted on positions 1 and 5 and joined by means of mP2 CLIPS to nucleate an α-helical structure. Native cysteines are replaced by cysteine-acm.

The binding of the antibody to each of the synthesized peptides was tested in a Pepscan-based ELISA. The peptide arrays were incubated with primary antibody solution (overnight at 4°C). After washing, the peptide arrays were incubated with a 1:1,000 dilution of an appropriate antibody peroxidase conjugate (anti-rabbit immunoglobulins HRP conjugate, DAKO, P0217) for one hour at 25°C. After washing, the peroxidase substrate 2,2’-azino-di-3-ethylbenzthiazoline sulfonate (ABTS) and 20 μL/mL of 3% H_2_O_2_ were added. After one hour, color development was measured. The color development was quantified with a charge coupled device (CCD) - camera and an image processing system. The values obtained from the CCD camera range from 0 to 3000 mAU, similar to a standard 96-well plate ELISA-reader.

## 3. Results

### 3.1. Ixochymostatin is a trypsin inhibitor-like (TIL) from *Ixodes scapularis*

The ixochymostatin has a protein-coding sequence of 91 amino acids, including a 19-amino acid signal peptide, with the mature sequence presenting a theoretical molecular weight of 7,882 Da and a pI of 5.00. The phylogenetic tree constructed using sequences of TILs from different species indicated that ixochymostatin and other TILs from *Ixodes* genus formed a separated clade (Figure 1A). Sequence and predicted structural analyses reveal the presence of one trypsin inhibitor-like (TIL) domain containing 10 conserved cysteine residues, which are predicted to form 5 disulfide bonds. The P1 residue, located in the inhibitory reactive site, is a tyrosine (Figure 1B). Alignment with other ticks and insect proteins belonging to the TIL family demonstrated conserved amino acid residues and similarity, with members of this clade sharing the tyrosine at the P1 position of the RCL (Figure 1A). The expression profile of ixochymostatin was assessed to evaluate its transcription throughout the tick-feeding process across various stages. Transcripts were detected in larvae, nymphs, as well as in the salivary glands and midguts of adult *I. scapularis* ticks (Figure 1C-F).

**Figure 1:**
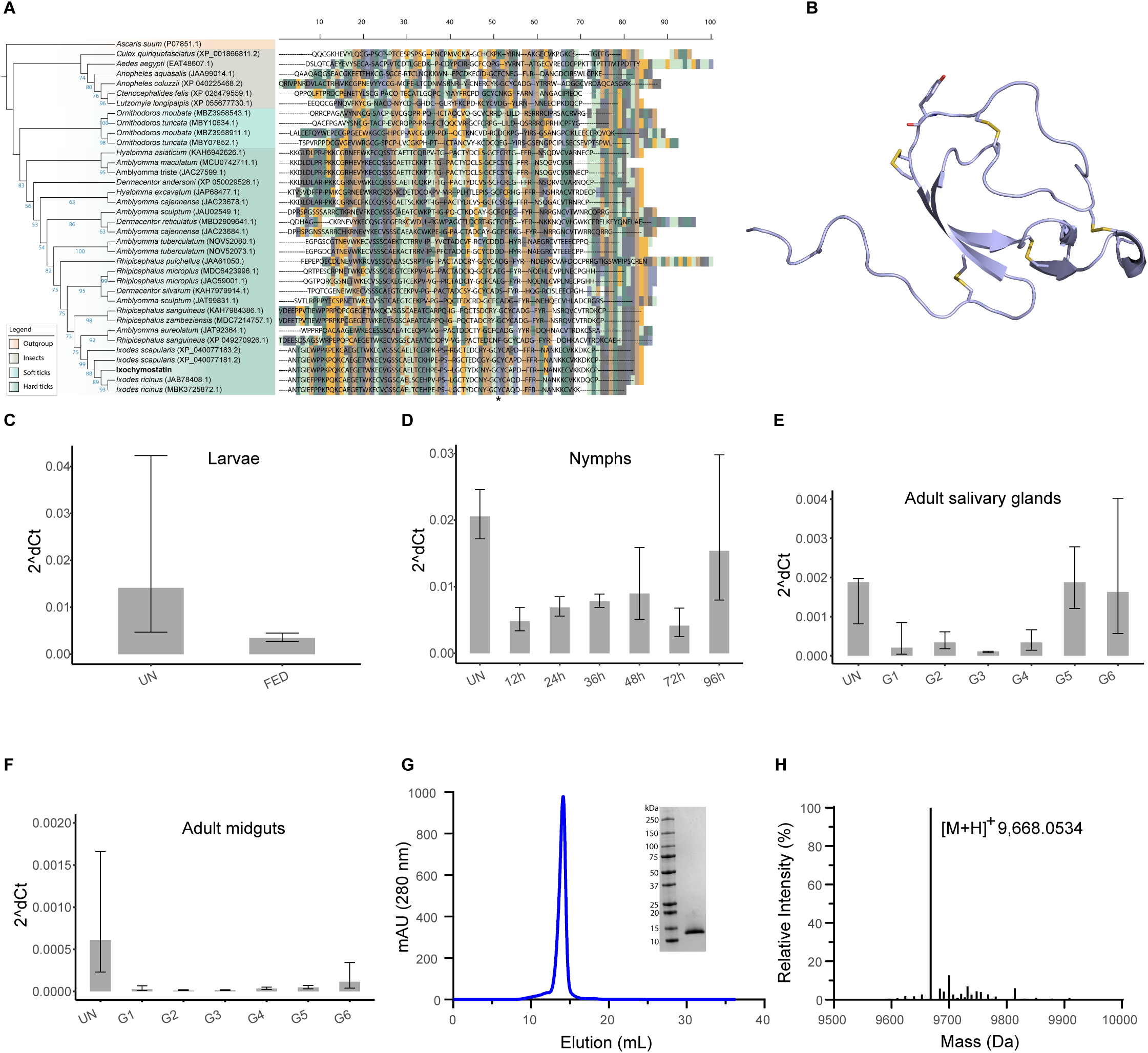
Ixochymostatin is a trypsin inhibitor-like (TIL) from *Ixodes scapularis*. (A) Maximum-likelihood tree reconstruction of trypsin inhibitor-like from different hematophagous arthropods. The tree was inferred using 1,000 replicates of ultrafast bootstrap (numbers at the nodes; values < 50 were omitted) and the substitution model WAG+I+G4. *Ascaris suum* (P07851) was used as an outgroup. GenBank accession numbers are in parentheses. Ixochymostatin is highlighted in bold, and an asterisk marks the P1 residue. (B) Structure of ixochymostatin predicted by AlphaFold2 and visualized using PyMol. Disulfide bridges are highlighted in yellow and the P1 residue (Tyr) is displayed as a stick. (C) qRT-PCR results showing the transcriptional profile of ixochymostatin in *Ixodes scapularis* unfed larvae (UN) and fed larvae (FED); (D) unfed nymphs (UN), 12h-fed nymphs (12h), 24h-fed nymphs (24h), 36h-fed nymphs (36h), 48h-fed nymphs (48h), 72h-fed nymphs (72h), and 96h-fed nymphs (96h); (E) salivary glands of unfed adults (UN), group G1 (G1), group G2 (G2), group G3 (G3), group G4 (G4), group G5 (G5), and group G6 (G6); and (F) midguts of unfed adults (UN), group G1 (G1), group G2 (G2), group G3 (G3), group G4 (G4), group G5 (G5), and group G6 (G6) [25]. (G) A chromatogram obtained from a Superdex 75 column run exhibits a single peak corresponding to purified ixochymostatin. The insert presents ixochymostatin solved in an SDS-PAGE 4-20% under reducing conditions, stained with Coomassie blue. (H) Intact mass of ixochymostatin performed by mass-spectrometry.

Recombinant ixochymostatin was successfully expressed in HEK293 cells and subsequently purified using affinity, ion-exchange, and gel-filtration chromatographies (Figure 1G). Protein homogeneity was confirmed by reducing 4-20% SDS-PAGE, revealing a single band (Figure 1G, inset), and by mass spectrometry, displaying a monomeric form with a mass of 9,668 Da (Figure 1H). The mass spectrometry data aligns with the mature sequence of ixochymostatin, followed by the 6x-his tag and enterokinase site present in the recombinant protein.

### 3.2. Ixochymostatin is a classical slow, tight-binding competitive inhibitor of chymotrypsin-like proteases

The inhibitory properties and specificity of ixochymostatin was verified in a screening assay against mammalian serine proteases related to host defense pathways. Incubation of ixochymostatin with each protease in a molar excess showed that ixochymostatin (1 µM) blocks partially or entirely the amidolytic activity of chymase, cathepsin G, and chymotrypsin (Figure 2A). Preincubation time between the protease and inhibitor significantly changes the progress curve’s behavior, suggesting that chymotrypsin inhibition by ixochymostatin occurs in a slow-binding mode (Figure 2B). Plotting the fractional velocity as a function of ixochymostatin at different protease concentrations resulted in directly proportional half-maximal inhibitory concentration values. We observed the higher the concentration of protease present, the higher the concentration of inhibitor required to reach half-maximal saturation of the inhibitor binding sites, suggesting ixochymostatin is a tight-binding inhibitor (Figure 2C). This tight-binding inhibition was confirmed by plotting the half-maximal inhibitory concentration values as a function of protease concentrations at a single fixed substrate concentration, yielding an apparent *K_i_* (*K_i_*^app^) of 16 nM (Figure 2D). Since IC_50_ usually does not properly reflect the real potency of tight-binding inhibitors, the Morrison equation [30] was used to fit fractional velocities at different inhibitor concentrations, allowing for the estimation of the *K_i_^app^* of 0.81 nM and a *K_i_* of 0.14 nM (Figure 2E). Moreover, measurement of ixochymostatin binding to a chymotrypsin-bound plasmon resonance surface (SPR) produced a dissociation constant (KD) of 7.6 × 10^−11^ M with an association rate constant (ka) of 1.6 × 10^5^ M−1s−1 and a dissociation rate constant (kd) of 1.2 × 10^-5^ s−1 (Figure 2F).

**Figure 2:**
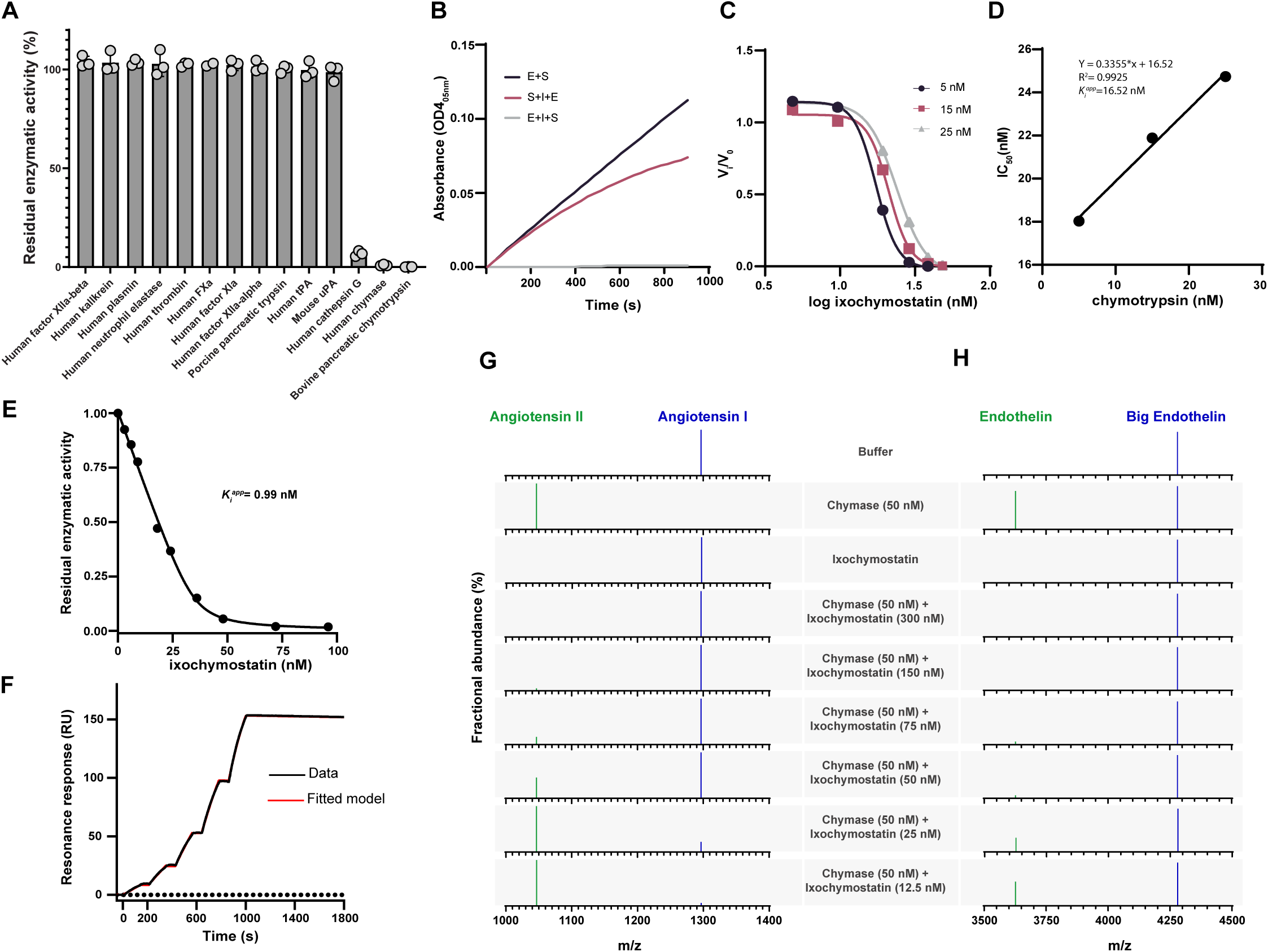
Ixochymostatin is a classical slow and tight-binding inhibitor of chymotrypsin-like proteases. (A) Inhibitory activity of ixochymostatin (1 μM) against different host-derived proteases. (B) Chymase (5 nM) hydrolysis of N-Succinyl-Ala-Ala-Pro-Phe-pNA (200 μM) was analyzed before (S+I+E) and after (E+I+S) previous incubation (10 min) with ixochymostatin (50 nM). Chymase (enzyme, denoted as E), ixochymostatin (inhibitor, denoted as I), and N-Succinyl-Ala-Ala-Pro-Phe-pNA (substrate, denoted as S). (C) Dose–response plot of fractional velocity (V_i_/V_o_) as a function of ixochymostatin (0–50 nM) at different chymotrypsin concentrations (5, 15, and 25 nM). (D) Linear behavior of IC_50_ values as a function of chymotrypsin concentration indicating a tight-binding mode of inhibition by ixochymostatin and its apparent *K_i_* (*K_i_*^app^). (E) Morrison’s plot of chymotrypsin (25 nM) residual activity as a function of ixochymostatin (0–96 nM) indicating a tight-binding competitive inhibition. (F) Surface plasmon resonance in the single cycle mode of chymotrypsin to ixochymostatin immobilized on the chip surface. Analyte concentrations were: 0 nM, 1.95 nM, 3.9 nM, 7.8 nM, 15.6, and 31.2 nM. The KD was determined by fitting experimental data to a 1:1 binding model (black line). Mass spectrometry results for (G) angiotensin II and (H) endothelin generation after incubation with chymase and ixochymostatin at different concentrations (12.5-300nM).

Since ixochymostatin has been shown to inhibit chymase, we tested whether it could modulate the conversion of natural substrates such as angiotensin I and big endothelin by chymase. Incubation with chymase converted angiotensin I (*m/z* 1296.68) to angiotensin II (*m/z* 1046.54), consistent with previous findings, by cleaving the C-terminal His–Leu dipeptide (Figure 2G). Chymase also converted big-endothelin (*m/z* 4280.92) to endothelin I (*m/z* 3626.587) at the Tyr– Gly bond (Figure 2H). Ixochymostatin dose-dependently inhibits the generation of angiotensin II (Figure 2G) and endothelin I (Figure 2H).

### 3.3. Ixochymostatin inhibits chymase and reduces vascular permeability induced by this protease

The integrity and permeability of the endothelial cell barrier at the interface of capillary vessels and tissues can be influenced and even compromised by proteases. Chymase increases the permeability of endothelial cells to FITC-dextran (Figure 3A) by disrupting VE-cadherin and cell-cell contacts (Figure 3B). Conversely, endothelial cells pretreated with ixochymostatin exhibit reduced permeability when exposed to chymase (Figure 3A) and were protected against chymase-induced VE-cadherin degradation, as confirmed by immunostaining (Figure 3B).

**Figure 3:**
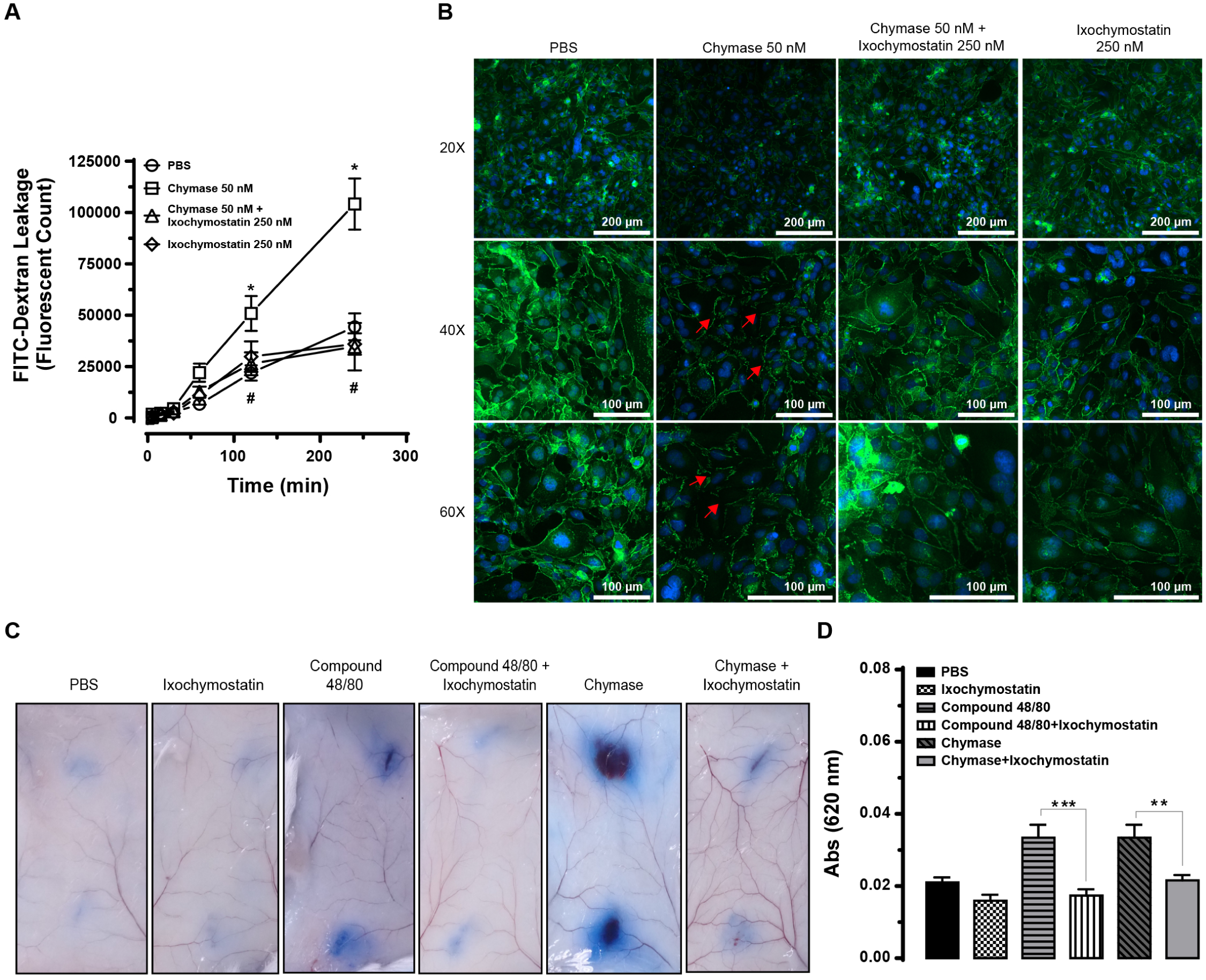
Ixochymostatin inhibits chymase and impairs vascular permeability *in vitro* and *in vivo*. (A) FITC-dextran assay was used to measure endothelial cell monolayer permeability on HMEC-1 cell line. The fluorescence intensity was quantified at different time points (0–4 h) after HMEC-1 treatment with phosphate-buffered saline, pH 7.4 (PBS), chymase, chymase plus ixochymostatin, and ixochymostatin (*p < 0.05 vs PBS; #p < 0.05 vs chymase 50 nM). (B) Immunostaining for VE-cadherin on HMEC-1 surface. Images are presented at different magnifications (20, 40, and 60 x). The red arrows indicated points of chymase-induced VE-cadherin degradation. (C) Vascular permeability to the Evans blue dye was evaluated *in vivo* using BALB/c mice. The following treatments were applied in two spots per animal: (i) phosphate-buffered saline pH 7.4 (PBS), (ii) compound 48/80 (1 µg), (iii) ixochymostatin (5 µg), (iv) chymase (3 µg), (v) compound 48/80 (1 µg) plus ixochymostatin (5 µg), and (vi) chymase (3 µg) plus ixochymostatin (5 µg). After 1 hour, the skin encompassing both injection sites was carefully removed and photographed. (D) Evans blue dye spots on the skin were excised, extracted with 50 % formamide (1.5 mL) and measured at OD_620nm_ (**p < 0.01 and ***p < 0.001).

To investigate if ixochymostatin was functional *in vivo*, the Miles assay in mice was employed. Intradermal injection of compound 48/80 (a mast cell degranulator) increases vascular permeability into the subcutaneous tissue, which was estimated by measuring Evans blue dye fluid extravasation in mouse skin. Compared to PBS injection, compound 48/80 significantly increased vascular permeability into the skin subcutaneous tissue. However, co-injection with ixochymostatin significantly reduced the amount of Evans blue extravasation. Similarly, intradermal injection of chymase increases vascular permeability into the subcutaneous tissue, and co-injection with ixochymostatin significantly reduces the amount of Evans blue extravasation (Figure 3C and D).

### 3.4. Antibodies against ixochymostatin neutralize its inhibitory functions, and potential epitopes responsible for this neutralization have been identified

We have investigated whether antibodies against ixochymostatin could neutralize its inhibitory functions. Monospecific antibodies neutralized in a dose-dependent manner the inhibitory function of ixochymostatin against chymotrypsin (Figure 4A). To determine putative epitopes responsible for this effect, ELISA was performed using purified IgG from pre-immune sera or from serum raised against the ixochymostatin. These were used against immobilized overlapping peptides from the full mature sequence of ixochymostatin to screen for epitopes, resulting in three epitope candidates. Several binding peaks were found with linear and conformational mapping. We observed three strong binding regions, which were not present in a pre-immune antibody (Figure 4B). The spatial location of the putative epitope binders in the tridimensional structure of ixochymostatin and in the complex with chymase was observed (Figures 4C and D). We have identified one epitope close to the interface between ixochymostatin and chymase. Antibody binding to this region could potentially be responsible for neutralizing the inhibitor activity, as the antibody binding would physically block the interaction site (Figure 4D).

**Figure 4:**
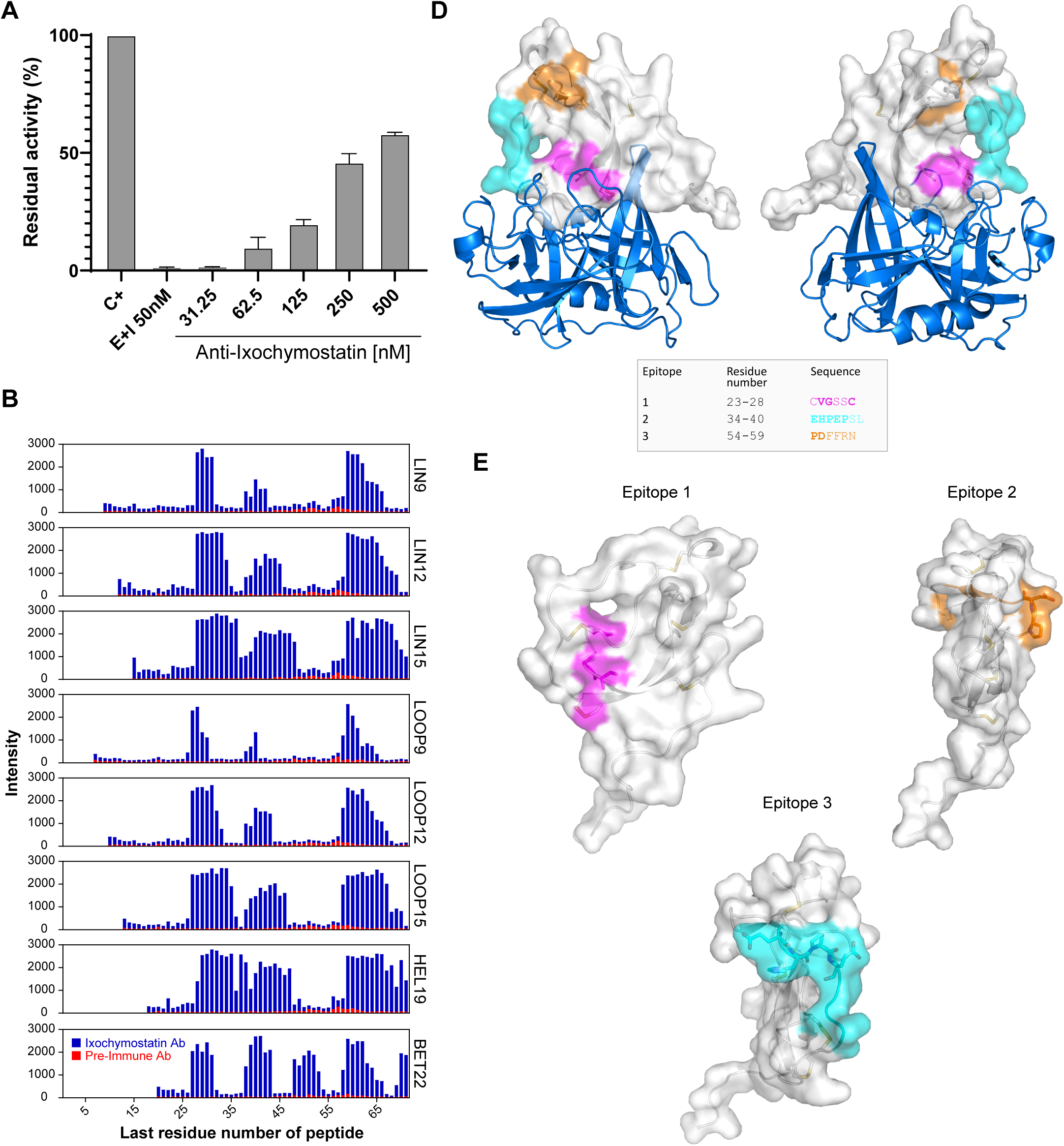
Antibodies against ixochymostatin neutralize its inhibitory properties. (A) Effect of antibodies upon ixochymostatin inhibitory activity. Monospecific antibodies (0 – 500 nM) were incubated with ixochymostatin (5 nM) at 37 °C for 30 min before the addition of chymotrypsin (5 nM), following a new 15-min incubation. Substrate was added and protease kinetics was monitored. The data was plotted as residual enzymatic activity (%) compared with the positive control (C+). (B) Overlay binding profiles of ixochymostatin recorded for monospecific antibodies on linear and conformational peptide arrays. ELISA intensities for anti-ixochymostatin (blue) and pre-immune control (red) were overlaid in a bar plot. Signal intensities are plotted on the y-axis and positions of the last residues of a peptide with respect to the target sequence are on the x-axis. LIN, linear; LOOP, looped; HEL, helical; BET, beta-turn. (C) Epitopes identified by the Pepscan analysis and their location in the predicted three-dimensional structure of ixochymostatin in complex with human chymase. (D) Epitopes identified by the Pepscan analysis and their location in the predicted three-dimensional structure of ixochymostatin.

## 4. Discussion

Ticks are obligate hematophagous ectoparasites capable of blood-feeding on a diverse variety of hosts, ranging from mammals to reptiles and amphibians [33]. Hard ticks typically exhibit natural behavior by feeding on a host for several days to weeks to complete their blood uptake successfully. Their saliva contains a rich cocktail of molecules displaying anti-inflammatory, anti-platelet aggregation, vasodilatory, anti-clotting, and immunomodulatory properties [2]. These salivary molecules can also play crucial roles in pathogen transmission simply by suppressing the host immune response at the tick-bite site [6]. Despite advancements in understanding tick sialomes, many salivary proteins still await characterization.

In ticks, serine protease inhibitors are one of the most prominent components identified in the sialotranscriptomes and sialoproteomes [5]. This family comprises several members, including serpins, Kunitz-type, Kazal-type, and trypsin inhibitor-like [13]. Although highly abundant in the saliva and salivary glands of ticks, data on the functional characterization of proteins belonging to the trypsin inhibitor-like (TIL) family are still scarce. In hematophagous arthropods, only a small number of proteins containing the TIL domain have been identified and described. Here, we present the first functionally characterized TIL from *Ixodes scapularis*. Ixochymostatin inhibits chymase, cathepsin G, and chymotrypsin. All of these proteases are categorized as chymotrypsin-like serine proteases. Ixochymostatin was identified as a slow and tight-binding inhibitor of chymotrypsin.

As ticks attach to the host’s skin to begin feeding, mast cells, neutrophils, and macrophages, among other immune cells, infiltrate the tick feeding site [34]. Cathepsin G is a serine protease that is released from neutrophil azurophilic granules after cell activation and has been implicated in the regulation of cell recruitment (through cytokine and chemokine processing), cell migration (through cleavage of adhesion molecules), and blood coagulation (through platelet aggregation induction via PAR4 activation) [35–38]. Similarly, chymase is the major protein stored and secreted by mast cells. Humans have one gene encoding chymase, while other mammal could have several chymases [39]. Thus, the types, amounts and properties of these serine peptidases vary by mast cell subtype, tissue, and mammal of origin. While most chymase-like proteases, such as the human chymase, primarily hydrolyze chymotryptic substrates [40], certain rodent chymases exhibit elastolytic activity due to mutations [41]. The predicted cleavage site P1–P1’ residue of ixochymostatin is the Tyr-Asp, which corresponds to the specific cleavage site for chymotrypsin-like proteases, which preferentially hydrolyses peptide bonds after aromatic amino acid residues [42]. The secretion of chymases contributes to inflammation, matrix degradation, vascular reactivity and tissue remodeling through various mechanisms. These include the degradation of pro-coagulant, matrix, growth, and differentiation factors, generation of vasoactive peptides, as well as the activation of proteinase-activated receptors, urokinase, and metalloproteinases. Additionally, chymases play a role in modulating immune responses by hydrolyzing and modifying chemokines and cytokines [43]. Notably, at least one chymase has been found to confer protection against intestinal worms in mice. Previous studies revealed that mice lacking mast cells were unable to build resistance against *Haemaphysalis longicornis* larvae [44]. Conversely, while wild-type mice exhibiting an intact mast cell response demonstrated resistance to tick feeding by triggering significant degranulation of these cells at the feeding site [45]. Thus, the inhibition of chymase and cathepsin G by ixochymostatin suggests this inhibitor may have a role in modulating inflammation, platelet aggregation and vascular responses during tick feeding [2].

In addition to the pro-inflammatory roles of mast cell chymase [43], secretion of chymase by mast cells can be responsible for the maturation of vasoactive peptides such as angiotensin II and endothelin, both of which play roles in regulating vascular blood flow and hemodynamic balance [46,47]. In this context, the inhibition of chymase by ixochymostatin may suggest a mechanism employed by ticks to hinder vasoconstriction induced locally at the skin by the generation of vasoactive peptides, thereby facilitating the flow of blood necessary for their meal acquisition.

Although there are no empirical data, the observed mast cell degranulation at the tick feeding site is expected to result in the release of chymase, which will lead to enhanced inflammatory and vascular responses [43]. In fact, ixochymostatin effectively protected the endothelial cell barrier against chymase-induced VE-cadherin degradation, strengthening cell-cell interactions. As a consequence, ixochymostatin significantly reduced the increase in endothelial cell permeability promoted by this protease. To provide a second measure of vascular leakage, supporting the effects observed in endothelial cells, the vascular permeability assay *in vivo* was performed by quantifying fluid extravasation in mouse skin using the Miles assay. Intradermal injection of compound 48/80, an agonist of mast cell degranulation, and even the injection of chymase itself increased vascular permeability into the subcutaneous tissue. Results showed that ixochymostatin significantly inhibited the permeability induced by compound 48/80 or chymase. Consequently, ixochymostatin emerges as a novel tick protein modulating vascular permeability, suggesting that *I. scapularis* ticks might utilize native ixochymostatin to dampen inflammation and vascular reactivity in response to tick feeding by inhibiting chymase secreted by degranulating mast cells at the tick feeding site. The presence of tick salivary proteins targeting cathepsin G and chymase has been described in other tick species [48,49], suggesting the importance of modulating these proteases for ticks to acquire a blood meal.

Considering the use of salivary proteins as potential antigens for tick control, the results obtained in this study are encouraging. Antibodies raised against ixochymostatin were able to neutralize its inhibitory properties. This result is important in the context of anti-tick vaccines, where immunization with this inhibitor would raise antibodies that can neutralize the functions of ixochymostatin at the feeding site during tick feeding, potentially impairing tick feeding and pathogen transmission. Pepscan analysis identified three putative epitopes for ixochymostatin. The predicted structure of the complex between ixochymostatin and a known target protease, chymase provides better insight into the region where ixochymostatin interacts with the protease. We have identified one epitope surrounding the interface of interaction between ixochymostatin and chymase. Antibody binding to this specific epitope could potentially be responsible for the observed neutralization activity, as the antibody binding would physically block the interaction site. Given the host’s defense modulatory effects of ixochymostatin and the assumption that they are crucial for the blood meal, neutralizing this protein could potentially have a negative impact on tick feeding and fitness. One approach to consider in future immunization studies with this protein is the use of epitope-based vaccines, which involves identification of immunodominant epitopes capable of inducing a protective immune response. Immunizing animals with a neutralizing peptide, instead of the entire protein, would help avoid the generation of non-protective antibodies, fostering a more targeted response against the antigenic determinants. Additionally, due to their small size, it would be possible to build formulations using multiple epitopes from the same or different proteins.

## 4. Conclusion

We have for the first time functionally characterized TIL from *Ixodes scapularis* ticks. Ixochymostatin inhibits chymase, cathepsin G, and chymotrypsin, and it was identified as a slow and tight-binding inhibitor of chymotrypsin. Inhibition of chymase effectively blocks vasoactive peptide generation and impairs endothelial cell and vascular permeability promoted by this protease. Consequently, ixochymostatin emerges as a novel tick protein modulating vascular responses. Antibodies generated against ixochymostatin neutralize its inhibitory properties. Through epitope mapping screening, we identified three main regions that might be responsible for this neutralization. This discovery enriches our understanding of the complex interactions between ticks and the vertebrate host, providing valuable insights for future research and potential applications in addressing tick-borne diseases.

## CRediT authorship contribution statement

Conceptualization: LAM, MB, LT; Investigation: LAM, MB, JK, SL, LCSP, BS, YZ, JFA, LT; Funding Acquisition: LT; Writing - Original Draft Preparation: LAM, LT; Writing – Review and Editing: LAM, MB, JK, LCSP, SL, BJS, YZ, JFA, LT.

## Acknowledgment

This work utilized the computational resources of the National Institutes of Health High-Performance Computing Services Biowulf cluster (http://hpc.nih.gov). LT was supported by the Division of Intramural Research of the National Institute of Allergy and Infectious Diseases (Z01 AI001337-01). MB was supported by funding from Conselho Nacional de Desenvolvimento Científico e Tecnológico, Ministério da Ciência e Tecnologia, Brazil (Chamada Universal MCTI/CNPq N° 10/2023, Grant 408915/2023-4) and Fundo de Incentivo à Pesquisa e Eventos (FIPE-HCPA, Grants N° 19-0001; 2024-0056; 2023-0258; 2023-0053) at Hospital de Clínicas de Porto Alegre.

